# Cell age drives asynchronous transcriptome aging

**DOI:** 10.1101/2023.05.31.543091

**Authors:** Ming Yang, Benjamin R. Harrison, Daniel E.L. Promislow

## Abstract

Organs age at different rates within a single individual. Such asynchrony in aging has been widely observed at multiple levels, from functional hallmarks, such as anatomical structures and physiological processes, to molecular endophenotypes, such as the transcriptome and metabolome. However, we lack a conceptual framework to understand why some components age faster than others. Just as demographic models explain why aging evolves, here we test the hypothesis that demographic differences among cell types, determined by cell-specific differences in turnover rate, can explain why the transcriptome shows signs of aging in some cell types but not others. Through analysis of mouse single-cell transcriptome data across diverse organs and ages, we find that cellular age explains a large proportion of the variation in the age-related increase in transcriptome variance. We further show that long-lived cells are characterized by relatively high expression of genes associated with proteostasis, and that the transcriptome of long-lived cells shows greater evolutionary constraint than short-lived cells. In contrast, in short-lived cell types the transcriptome is enriched for genes associated with DNA repair. Based on these observations, we develop a novel heuristic model that explains how and why aging rates differ among cell types.

## Introduction

In his poem The Deacon’s Masterpiece, Oliver Wendell Holmes described a carriage so artfully constructed that it ran flawlessly for 100 years, at which time every single part failed simultaneously (Holmes 1923). The real world of organismal aging is, of course, very different. As animals age, different organs and functions fail at different rates and in different ways (Rando and Wyss-Coray 2021). In humans, hair color, reproductive capacity, and heart function all decline faster than cognitive and gastrointestinal function (Khan et al. 2017). Heterogeneity in aging among organs is observed not only in terms of structural and functional changes, but also at the molecular level. For example, in mice, the timing and degree of age-related change in the transcriptome differ among organs (Schaum et al. 2020). Similar among-organ differences are seen in the aging of the proteome (Ori et al. 2015). However, these observations are based on analysis of bulk tissue, which masks variation among cells within tissues. Analysis of aging in single-celled organisms has established transcriptome-wide effects of cellular age, including declining coordination among genes expressed within cells, and increased discordance in gene expression profiles among cells (Rando and Wyss-Coray 2021).

Studies using single-cell transcriptomics have made substantial contributions to our understanding of variation in aging among organs and cells. In particular, through analysis of single-cell transcriptome data among cell types within and across organs, researchers have found strong evidence of cell-specific differences in the way that age impacts gene expression levels, gene expression variability, and cell-to-cell heterogeneity (Martinez-Jimenez et al. 2017; Kimmel et al. 2019; Angelidis et al. 2019; Salzer et al. 2018; Ximerakis et al. 2019; Enge et al. 2017; Zhang et al. 2021; Ibañez-Solé et al. 2022; Bahar et al. 2006). For example, in mice, age-related changes in gene expression occur in natural killer cells in the lung and spleen, but to a much less extent in B cells from the same organ (Kimmel et al. 2019). Although the relationship between asynchronous variation in the transcriptome and functional decline among organs is speculative, this variation among cell types may, in turn, help us to understand variation in rates of aging among tissues and organs (Cohen et al. 2022).

Despite the potential for single-cell analysis to uncover the asynchrony in tissue and organ aging, we lack a compelling explanation for why transcriptome variation changes with age in some organs but not in others. Prior studies have proposed that transcriptome variation builds from age-driven ‘noise’ in gene expression, which may be caused by somatic mutation (Enge et al. 2017), epigenetic drift (Fraga et al. 2005), or other damage to gene expression programs (Levy et al. 2020; Warren et al. 2007). Others have argued that properly regulated age-dependent gene expression, some of which may prevent or repair cellular damage, is an important component as well (Perez-Gomez et al. 2020; Ibañez-Solé et al. 2022). Some of these potentially causal processes are known to differ among cell types (Enge et al. 2017; Moore et al. 2021; Li et al. 2021). But the root cause of this variation might actually be related to the underlying population dynamics of different cell types. In particular, Warren *et al*. suggested that asynchrony in transcriptome variation, rather than being a direct reflection of organ aging, might instead be explained by differences in cell turnover rates (Warren et al. 2007). Cells from slowly-renewing tissues are more likely to have cellular ages similar to the organism’s age, while fast-turnover cells, being constantly replenished, stay relatively young throughout the life of an organism. This explanation is consistent with the observation that age-related increases in transcriptome variability have been detected in low-turnover cardiomyocytes (Bahar et al. 2006), but not in high-turnover granulocytes, naive B and T cells, or endothelial cells of the lung (Warren et al. 2007; Ibañez-Solé et al. 2022; Kimmel et al. 2019).

Cell turnover rate varies by orders of magnitude among cell types, from fast turnover rates in blood or gut epithelial cells, which are renewed every day (Sender and Milo 2021; Richardson et al. 2014), to very slow turnover in neurons, whose renewal during an individual’s lifetime might only occur in specific regions of the brain, if at all (Spalding et al. 2013). The distribution of ages in a population of cells of a given type will depend on cell turnover rate and the age of the organism, and with a few simplifying assumptions, we can thus estimate the age distribution of a population of cells of a given type. Using these age estimates, here we show that cell-type specific differences in cellular age account for a substantial proportion of the variance in how age affects cell-to-cell variation in the transcriptome. This effect appears to supersede the role of niche identity on aging of post-mitotic cells. We also find that long lived cells tend to express genes associated with proteostasis and mitophagy, and that the transcriptome of long-lived cells shows signs of stronger selective constraint relative to genes expressed in short-lived cells. Short-lived cells, in contrast, show higher expression of genes associated with DNA repair. We suggest that the function of short-lived cells may be optimized by evolutionary forces of selection and a somatic version of Williams’ Antagonistic Pleiotropy phenomenon (Williams 1957). Taken together, these results identify a critical effect of cellular age on the transcriptome, and suggest a heuristic model for the different strategies that short-lived and long-lived cells employ to ensure cell function over the course of an organism’s life.

## Results

### Cell turnover explains asynchronous trajectories of transcriptome variation with age

To investigate the relationship between transcriptome variation and cellular age across different cell types, we obtained cell type-specific cellular lifespan data in rodents from Sender and Milo (2021) and integrated it with single cell transcriptome data from the mouse aging atlas, *Tabula Muris Senis* (TMS) (Zhang et al. 2021). Together, this yielded 39 combinations of mostly post-mitotic organ:cell types, within their tissue context, with both lifespan estimates and age-specific transcriptome profiles (**Fig. 1A, Methods, Supplemental Table S1**). We used published measures of cell lifespan (*T*) to estimate the mean age of each cell type in young (3-month-old) and old (24-month-old) mice, respectively. Making the simplifying assumption that cell turnover rates are constant with age, we can estimate cell turnover rate *b* = 1 - *e*^-1/T^ (**Methods, Supplemental Methods**). Given that a cell cannot be older than the organism in which it is found, the distribution of cellular age among a given type will be a truncated exponential function determined by cell turnover rate and organismal age. We can then use organismal age to calculate the mean age of a population of cells with turnover *b*. This cellular demography model illustrates the degree to which cellular age increases with organismal age. For cell types with relatively slow cellular turnover, the mean cellular age increases concomitantly with organismal age. At the extreme, a cell that has no turnover will always be the same age as the organism in which it is found. In contrast, cells with very fast turnover will never be more than a few days old and, barring any changes in turnover rate with host age, the age distribution for that cell type will be constant throughout the life of the organism (**Fig. 1B, Supplemental Figs. S1-2, Supplemental Methods**).

**Figure 1.**
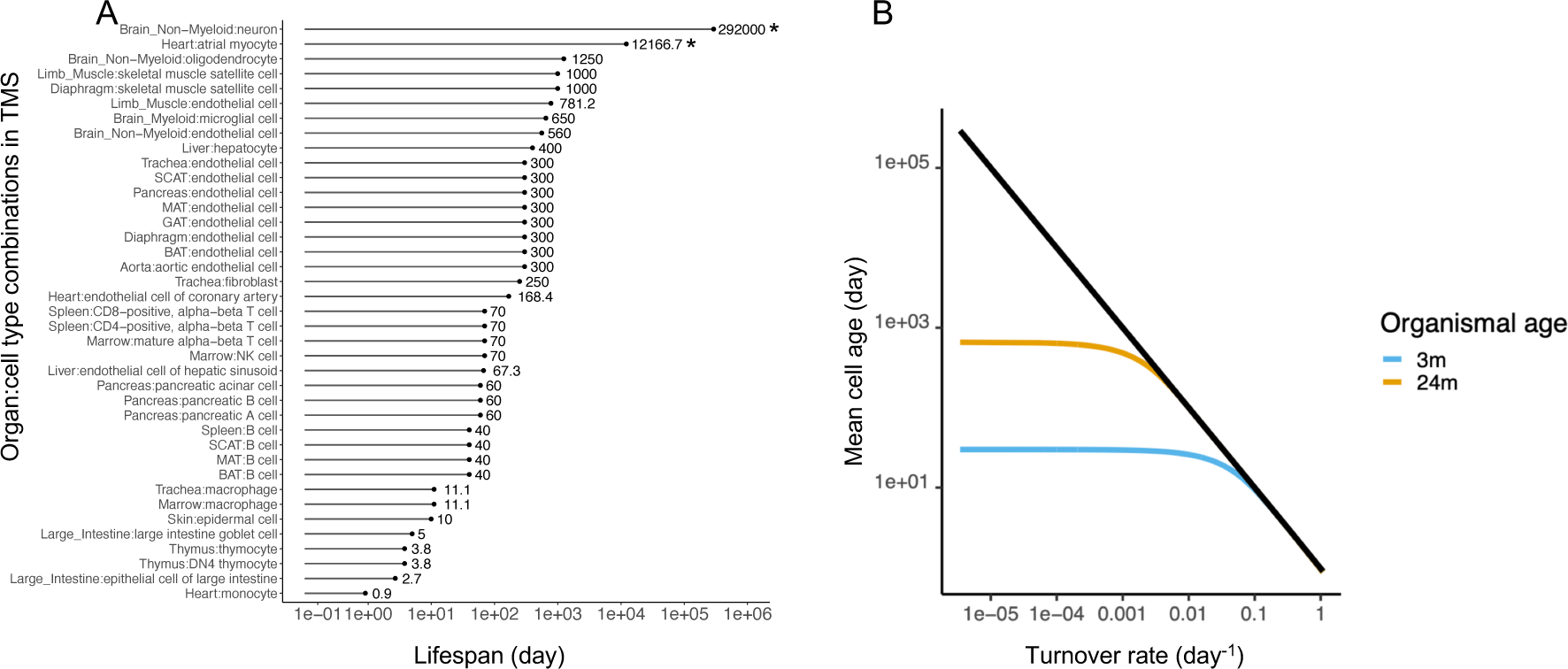
The distribution of cell lifespan data used in this study and of mean cell age distributions during organismal aging. **(A)** Summary of cellular lifespan data used in this study. The cellular lifespan data in rodents (Sender and Milo 2021) were aligned with cell types in the TMS data (Zhang et al. 2021) (see **Supplemental Table S1**). Note that in two cases, the statistical model used in Sender and Milo (2021) gives lifespan estimates that exceed the maximum lifespan of laboratory mice, highlighted with * for neuron and atrial myocyte. However, these values lead to minimal effects on the mean cell age during organismal aging. **(B)** Mean cell age as a function of cell turnover rate. The black line denotes the theoretical expectation for a cell with an exponential survival function in an infinitely long-lived organism. More realistically, we assume that cell age follows a truncated exponential distribution, such that the mean cell age is constrained by organismal age. We show mean cell age as a function of cell turnover rate for a 3-month-old mouse (blue) and a 24-month-old mouse (yellow), using the model in Equation 8 (**Supplemental Methods**).

To quantify cell type-specific transcriptome variability with the TMS data, we measured the variability of individual genes among cells of a given organ:cell type combination at young and old age separately. We then fitted a mixed model to estimate the effect of age on transcriptome-wide variability (β_age_, **Methods**). To minimize any confounding effects of a mean-variance correlation on gene expression (Eling et al. 2018), we limited our analysis to the 16% to 83% of genes that are not differentially expressed with age within each of the 39 organ:cell type combinations (P > 0.1, **Methods**) (Martinez-Jimenez et al. 2017). We found that gene-level variance increased with age in 35 of 39 organ:cell type combinations (β_age_ > 0, FDR < 0.001, **Supplemental Fig. S3**). Thus, most cell types show at least some increase in transcriptome variability with organismal age, consistent with previous work (Bahar et al. 2006; Kimmel et al. 2019; Angelidis et al. 2019; Enge et al. 2017; Luo et al. 2022; Wang et al. 2020).

Considering that slow turnover cells are calculated to show a more dramatic increase in cellular age between young and old animals compared to fast turnover cells (**Fig. 1B**), we hypothesized that such a shift in cellular age would correspond to larger changes in transcriptome variation. To test this prediction, we first calculated the increase in the mean cell age of each cell type from 3-month-old to 24-month-old mice (Δ_age_; **Supplemental Methods, Eq. 9**). We then examined the relationship between β_age_ and Δ_age_ across all cell types.

Cell lifespan estimates were derived for distinct cell types by Sender and Milo (2021), and in many cases a given cell type is distributed across different organs (e.g., we have TMS data for B cells found in brown adipose, mesenteric adipose, and subcutaneous adipose tissue). Tissue microenvironment may affect cell turnover rate, and there are organ-specific turnover estimates for 14 of the 39 organ:cell type combinations (**Methods**). The remaining 25 organ:cell type combinations correspond to 7 cell types, all without organ-specific turnover estimates. To avoid pseudoreplication we made the simplifying assumption that when organ-specific turnover was not known, β_age_ is best estimated at the cell type level by taking its mean within that cell type. This approach gave 21 independent estimates of both β_age_ and Δ_age_.

We found a striking positive correlation between cellular age and the effect of age on the transcriptome, with more than half of the variation in β_age_ explained by Δ_age_ (Pearson’s *r* = 0.714; P = 2.8 × 10^-4^) (**Fig. 2, Supplemental Table S2**). To confirm that this finding did not depend on our metric of transcriptome variability, we applied two alternative metrics (Angelidis et al. 2019; Levy et al. 2020) and found the same pattern (**Methods, Supplemental Figs. S4-6, Supplemental Table S2**).

**Figure 2.**
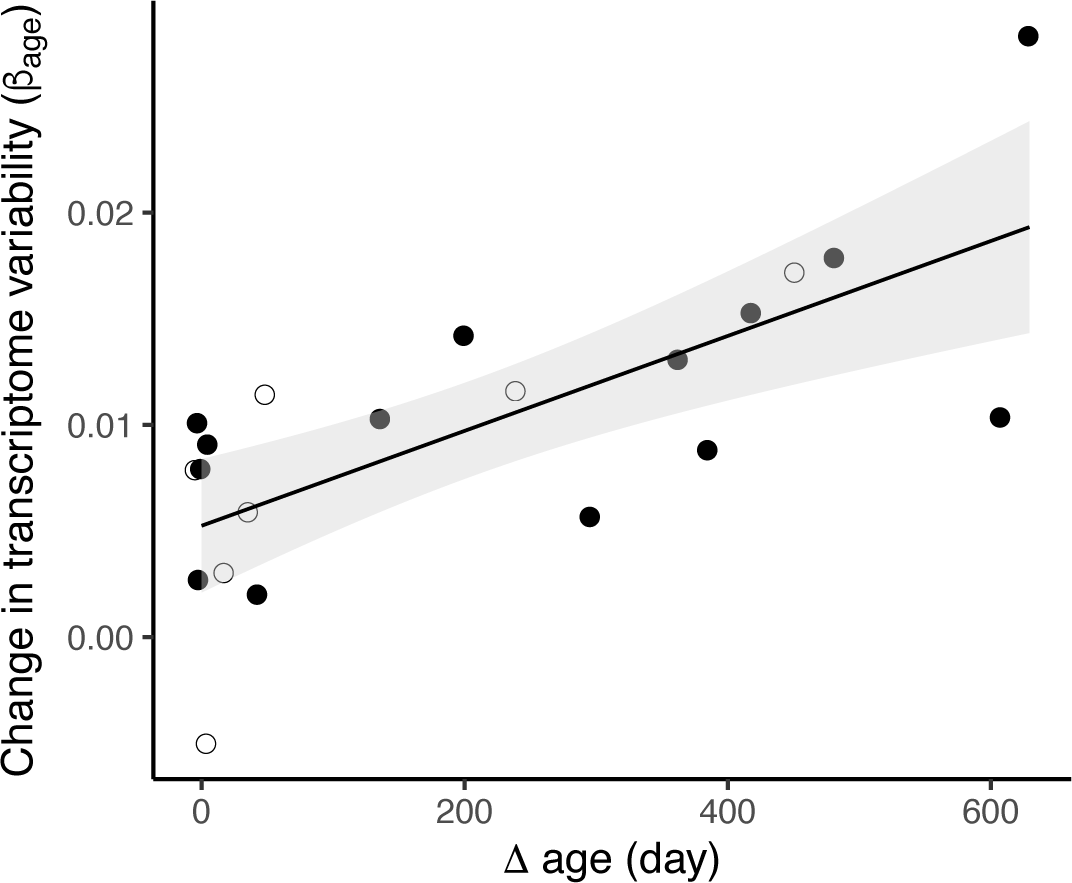
Cellular age dynamics predict the increase in transcriptome variability in aging mice. The mean change in transcriptome variability (β_age_, **Methods**) observed in cells over a 21-month period, from 24-month-old mice compared to cells from 3-month-old mice, is correlated with the change in estimated mean cell age for 21 cell types (Pearson’s r = 0.714, P = 2.8×10^-4^). The black line shows an ordinary least squares regression, with shading indicating 95% confidence intervals. Filled circles indicate the 14 cell types with cell- and organ-specific lifespan data, and open circles the 7 cell types without organ-specific lifespan estimates, and so their β_age_ is the mean from all organs in which they are detected (see **Methods**).

To explore the effect of tissue microenvironment on transcriptome variation, we separately considered organ-specific β_age_ for the instances where transcriptome data were available for specific cell types from more than one organ (**Supplemental Fig. S7**). We found that β_age_ appears to vary by organ (**Supplemental Fig. S7A**). However, this variation is likely driven by effects of tissue microenvironment on cell turnover. For example, in the case of endothelial cells, where we had estimates for both organ-specific turnover rate and organ-specific β_age_, variation in β_age_ is almost entirely explained by organ-specific Δ_age_ (**Supplemental Fig. S7B**). Taken together, our results demonstrate that a substantial proportion of the age-related changes in the gene expression variation among cell types can be explained by cellular age.

### Cell lifespan associates with signatures of selection and proteostasis

We next sought to explain why cells with less turnover and longer lifespan exhibit greater signs of aging. Given the deleterious and cumulative effects of somatic mutation, misfolded proteins, and dysfunctional mitochondria as organisms age (López-Otín et al. 2013), we asked if long-lived cells show increased expression of repair or maintenance processes in their transcriptome. Using Gene Ontology (GO) terms related to DNA repair, proteostasis, and mitophagy, we calculated the expression of each GO term in each cell, and the mean of these values for all cells in a cell type (**Methods**).

Consistent with our hypothesis, in cells of young mice we found that expression of three out of the six GO terms associated with chaperone-mediated processes, including protein folding, protein complex assembly, and autophagy, were positively correlated with cell lifespan, as was ‘autophagy of mitochondria’ (absolute Kendall’s τ > 0.34, FDR < 0.05, **Fig. 3A**). However, in cells from old mice, the relationship between cell lifespan and the expression of the chaperone-mediated terms was no longer significant (**Supplemental Fig. S9**). Counter to our expectation, expression of two of the six GO terms associated with DNA repair, while not associated with cell lifespan at young age, were *negatively* associated with cell lifespan at old age (absolute Kendall’s τ > 0.41, FDR < 0.05, **Fig. 3B**). That is, in older mice, long lived cells are less likely than short lived cells to express genes involved in DNA repair.

**Figure 3.**
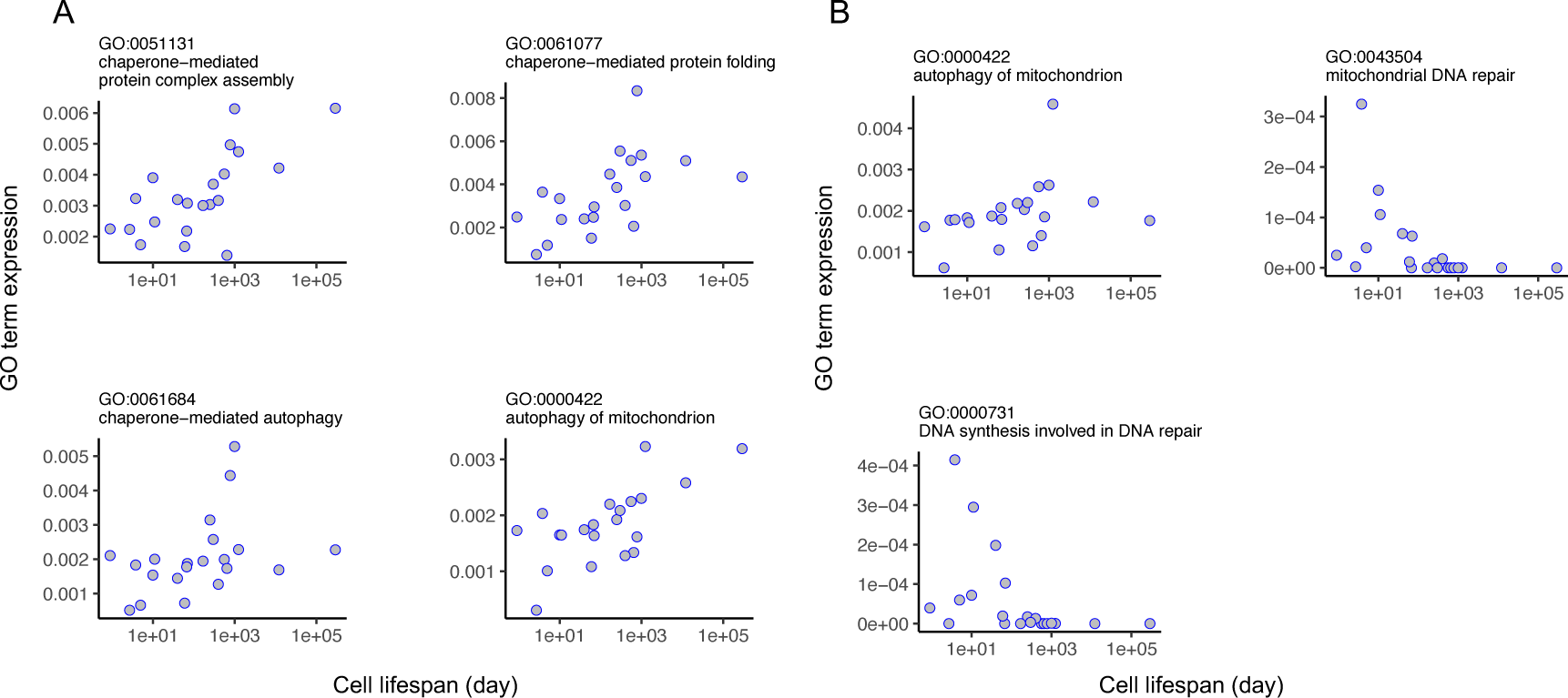
Cell lifespan correlates with chaperone mediated processes, mitochondrial autophagy and DNA repair in mice. The mean expression of genes in each GO term in 21 cell types at young age (**A**), and old age (**B**) correlates with the lifespan of those cell types. For all plots, the absolute Kendall’s τ > 0.34 (FDR < 0.05).

Our results point to the importance in long-lived cells of maintaining proteostasis through chaperone expression (Labbadia and Morimoto 2015). However, proteins vary in their dependence on chaperones. As previous works has shown, chaperone clients tend to be less stable than proteins that are not chaperone clients, and they carry higher levels of non-synonymous codon variation (*d*_N_), indicating less evolutionary constraint (Taipale et al. 2012). In light of these findings, we might expect that long-lived cells decrease their proteostatic burden by expressing proteins that are, on average, more stable, with relatively low *d*_N_/*d*_S_ ratios (Drummond and Wilke 2008; Sikosek and Chan 2014). Consistent with this expectation, we found that *d*_N_/*d*_S_ ratio in the expression profiles of cells from young and old mice was negatively correlated with cell lifespan (Pearsons’s *r* < -0.59, P < 5.0 × 10^-3^, **Fig. 4**). This result remained unchanged after removing the 76 genes that were more likely to be under recurrent positive selection (*d*_N_/*d*_S_ > 1, **Supplemental Fig. S10**) (Kryazhimskiy and Plotkin 2008). In addition, we found that the association became stronger as we limited the analysis to genes whose expression patterns were cell type-specific (**Fig. 4C, Methods**).

**Figure 4.**
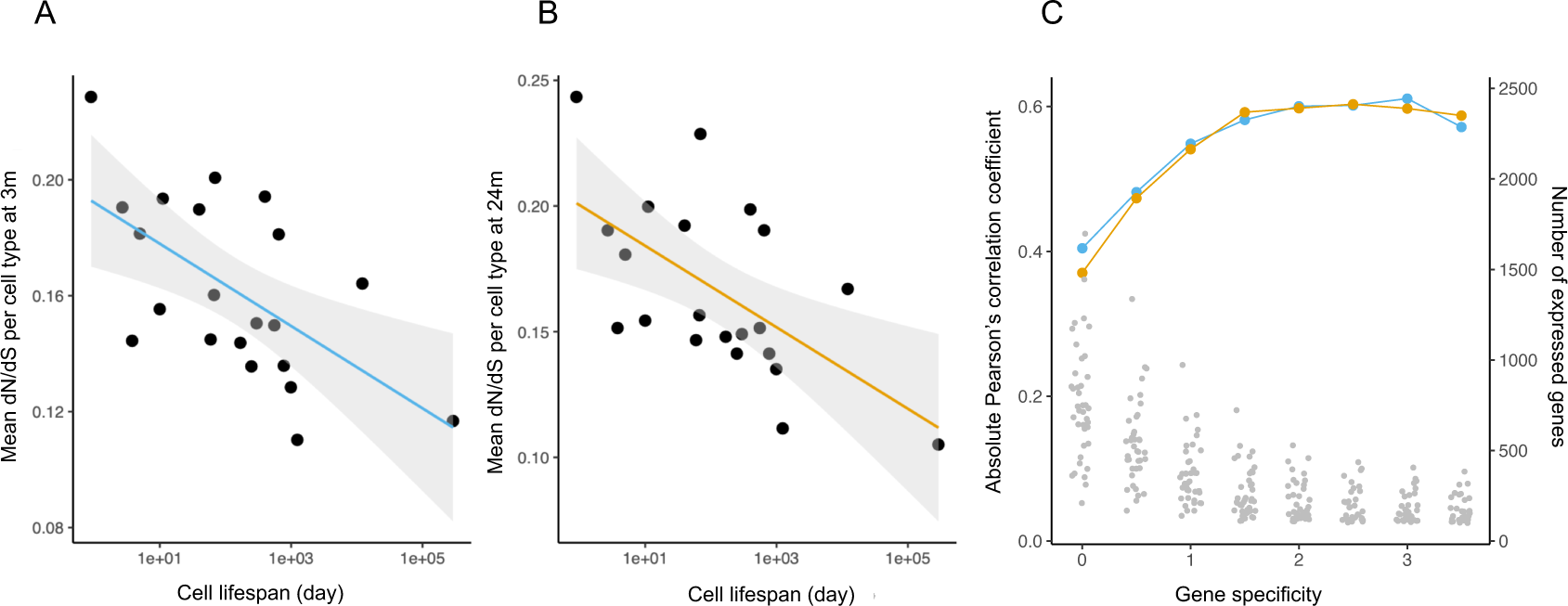
Cell lifespan predicts evolutionary constraint among expressed genes. The relaxation of evolutionary constraint (*d*_N_/*d*_S_) of genes expressed in 21 cell types in young (**A**) and old (**B**) mice is negatively correlated with cell lifespan (Pearson’s *r* = -0.6, P = 3.9 × 10^-3^, and Pearson’s *r* = -0.598, P = 4.2 × 10^-3^, for young and old mice, respectively). The median *d*_N_/*d*_S_ is measured per cell, using cell type-specific genes (median *d*_N_/*d*_S_ of genes above specificity score > 2, n = 116 to 971 genes, see **Methods**). The average *d*_N_/*d*_S_ of a cell type is the mean of *d*_N_/*d*_S_ across cells (n = 56 to 2251 cells, see **Methods**). The line in each figure shows an ordinary least squares regression with shading of 95% confidence intervals. **(C)** Correlation between dN/dS and cell lifespan for transcriptomes of increasing cell-type specificity (see **Methods**). The blue or yellow dots corresponding to the left-side axis show the correlation strength. The grey dots correspond to the right-side axis and show the number of expressed genes per cell type at each level of specificity.

## Discussion

It has long been recognized that the dynamics of aging—its onset, rate, and magnitude—vary across species (Gorbunova et al. 2014; Promislow 1991), and among individuals within species (Kirkwood 2005). But it is also clear that even within individual organisms, different structures and functions age at different rates. Surprisingly little work has been done to describe this variation (but see, for example, Hayward et al. 2015), and we lack a clear framework to explain why some elements age faster than others. A recent evolutionary model posits that those systems under stronger selection—that is, systems that contribute more to organismal fitness—should age faster than others (Moorad and Ravindran 2022), but the theory has yet to be tested, requiring estimates of the age-specific marginal effects on fitness of different components within an organism. Based on observations of cell-specific transcriptome variation with age in just four cell types, Warren *et al*. (2007) hypothesized that the degree to which the transcriptome shows signs of aging in a particular cell type is positively associated with the life expectancy of that cell type.

Here we provide the first comprehensive test of Warren *et al*.’s (2007) hypothesis. We show that in mice, the amount of age-related increase in transcriptome variation is indeed greater in long lived cells, such as neurons and muscle cells, relative to short-lived cells like monocytes and gut epithelial cells. While the functional consequence of gene expression variability remains an open question, it appears that differences in aging among cell types, as measured by transcriptome variation among cells within each type, are driven in part by cellular age. Moreover, we show that expression profiles of longer-lived cells are enriched for pathways involved in homeostasis, including chaperone-mediated processes and mitochondrial turnover. We cannot be sure whether higher expression of chaperone processes or mitophagy support long cellular lifespan, or if longer lifespan leads to greater opportunity for protein damage or mitochondrial malfunction, thereby inducing expression of genes associated with proteostasis or mitophagy. In support of the former interpretation, the proteins expressed in longer-lived cells tend to have evolved under stronger selective constraint.

To further understand how and why different cell types age differently, we need to consider the extrinsic and intrinsic factors that influence cell aging. Extrinsic factors that shape somatic cell lifespan likely include aging of stem cells (Jones and Rando 2011) and age-related changes in the cellular niche (Brunet et al. 2022). Intrinsic factors include, among other forces, somatic mutation, changes to epigenetic structure, oxidative and other damage, loss of mitochondrial function, diminution of homeostatic mechanisms, and apoptotic cell death (Elliott and Ravichandran 2016; Gladyshev et al. 2021). We consider each factor in light of our results.

First, there is extensive support for aging of stem cells, and for the hypothesis that variation in stem cell dynamics can influence traits associated with aging, especially cancer (Jones and Rando 2011; Tomasetti and Vogelstein 2015). However, our results fail to support a major role for stem cell aging as a driver of transcriptome variation among short-versus long-lived cells. Instead, we see the greatest age-related increase in transcriptome variation among cells with the slowest turnover, where there is relatively little stem cell activity, contrary to what one might expect if stem cell aging drives transcriptome variation.

Second, age-related changes occur in structure and function of the tissue microenvironment, or cellular niche (Brunet et al. 2022). How the niche affects cellular lifespan and transcriptome variation across an organism is unknown, but in each instance where a niche-specific cellular age is available in our study, it is strongly predictive of the effect of age on transcriptome variation (**Supplemental Fig. S7B)**. We speculate that in cells of a given type that occur in diverse organs, variation in rates of aging across different niches is driven by the effect of the niche on rates of cellular turnover, rather than directly on rates of increase in transcriptome variation.

Third, factors intrinsic to the cell, including somatic mutation, damage to other biomolecules, repair and cellular homeostatic mechanisms, and cell death, have all been implicated in aging (Gladyshev et al. 2021). Little is known about the degree to which cells differ in their baseline rates of certain kinds of damage, such as free radical damage to biomolecules. However, single cell genomic analyses have found significant heterogeneity in somatic mutation rates across cell types and organs, in both proliferating cells (e.g., Martincorena et al. 2018) and post-mitotic cells (Moore et al. 2021; Li et al. 2021). Somatic mutation, other damage to DNA, and age-related changes to the epigenome, are all associated with transcriptional variation, and so each may contribute as drivers of the transcriptional variation that accompanies cellular aging (Bozukova et al. 2022; Gyenis et al. 2023). We further note that mutation, DNA damage, and epigenetic change need not directly affect the expression of genes in cis, but rather any indirect effects on gene regulatory mechanisms or cell state may be sufficient to drive transcriptional variation within a cell type (Schumacher et al. 2021).

Additionally, programmed cell death (PCD), triggered by cellular damage, directly contributes to cell turnover. This creates opportunities for selection within populations of cells and should have effects on the transcriptome. Programmed cell death is critical to maintaining tissue function and preventing cancer, and so has received a great deal of attention (Pistritto et al. 2016). And yet, due to the complexity and variety of apoptotic mechanisms and their associated biomarkers, we lack a comprehensive picture of the rates and regulation of apoptotic cell death across cell types (Elliott and Ravichandran 2016). We further discuss the role of cell death in a model of cell turnover and organismal aging below.

### A Heuristic Model

We found compelling evidence that cells with relatively slow turnover rates have much greater age-related increase in transcriptome variance, and identified additional correlates that point to potential mechanisms that could account for this pattern. In **Figure 5** we present a heuristic model to help us integrate our findings, and to reconcile observations that might seem, on the face of it, to be contradictory.

**Figure 5.**
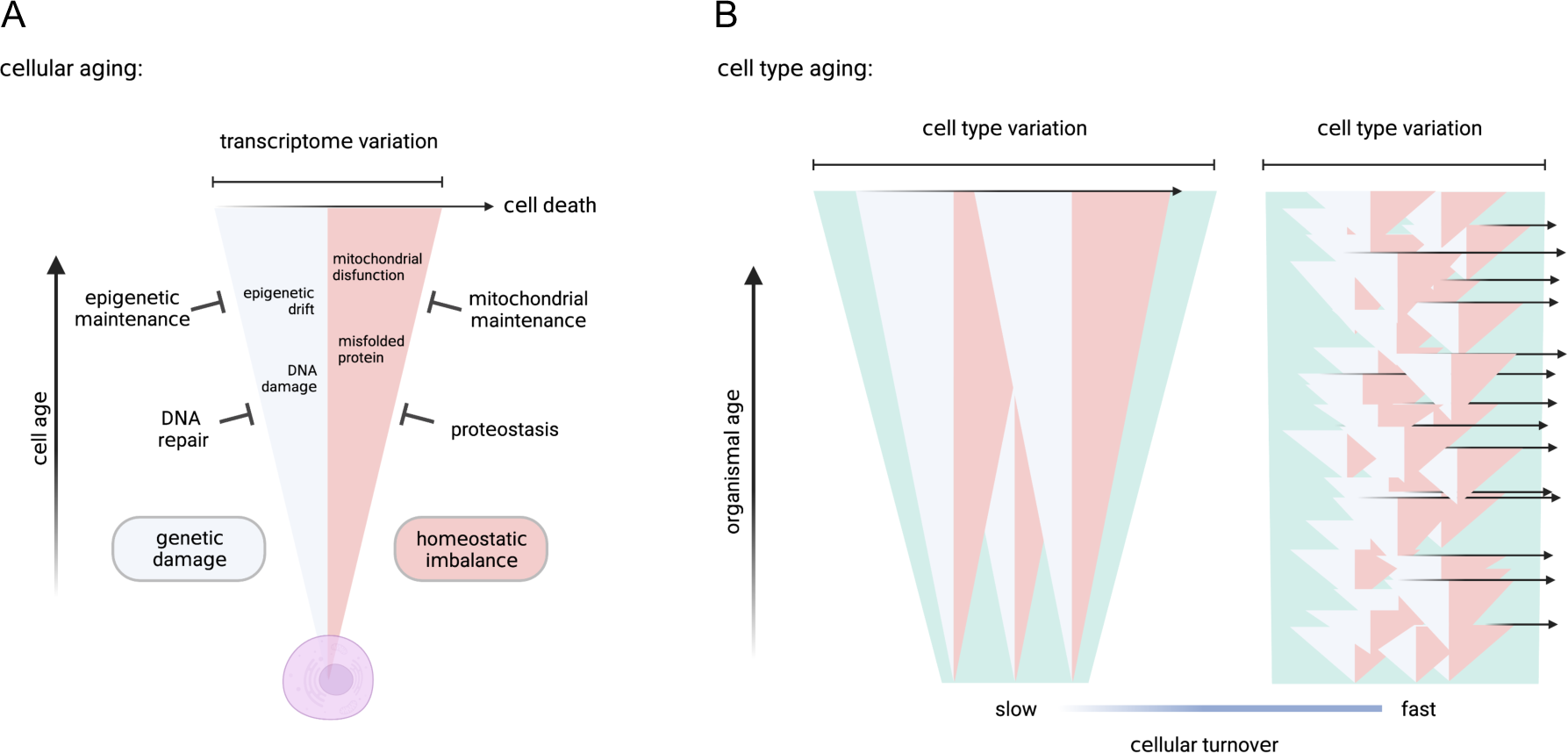
Models of cellular aging and cell type-specific transcriptome variation. Here we present a model depicting the accumulation of transcriptome variation with cellular age. **(A**) As a cell ages, it accumulates genetic damage (light blue) and cellular homeostasis becomes increasingly imbalanced (pink), either of which can directly or indirectly increase transcriptome variation. The rates of accumulation of damage and loss of homeostasis are counteracted by repair and maintenance processes, which in turn slow the rate of increase in transcriptome variation. As cellular transcriptome variation increases, so does the likelihood of triggering cell death (rightward arrow) (Tower 2015). (**B**) The model of cellular aging when applied to populations of long-lived cells (slow turnover) and short-lived cells (fast turnover) describes the accumulation of transcriptome variation among a cell type (green). Long-lived cells age together with the organism, and transcriptome variation accumulates among these cells as the organism ages. In short-lived cells, damaged cells are frequently replaced by new cells, which prevents accumulation of variation at the cell type level.

As cells age, we might expect deleterious changes in the transcriptome due to direct alteration of DNA, including somatic mutation and epigenetic drift (Bahar et al. 2006; Schumacher et al. 2021), which are counteracted or avoided by repair and maintenance processes (Yousefzadeh et al. 2021; Gyenis et al. 2023). Changes to the transcriptome could also be due to improper gene regulation because of loss of cellular homeostasis, and cells have evolved elaborate systems to maintain homeostasis through protein chaperones, autophagic recycling of cell components, and so forth (Santra et al. 2019; Lima et al. 2022).

In cells, all of these factors—mutation, oxidative damage, heat stress, etc.—will lead to the gradual accumulation of variation in the transcriptome with age (**Fig. 5A**). Increased expression of genes associated with proteostasis and mitophagy, and expression of proteins with relatively high stability, appear to be ways that long-lived cells mitigate this damage.

Notably, we observe that the expression levels of genes associated with DNA repair are highest not in the longer-lived cells, but in short-lived ones. While this observation might seem to contradict our model, we believe it is the exception that proves the rule. Many different factors lead to DNA mutation and damage, and mitotic division plays a major role (Tomasetti and Vogelstein 2015). Cells with little turnover are protected from this source of damage. While fully differentiated short-lived cells have little time to accumulate DNA errors, they could inherit DNA damage passed down by the long-lived progenitor stem cells from which they are derived.

When considering these various sources of accumulated error and damage, we are not suggesting that short-lived cells are immune to their effects. In fact, it is likely that short lifespan goes hand in hand with a much higher rate of accumulation of damage and failure. However, rapid turnover and replacement of these cells means that most cells in a population of short-lived cells are very young, with little damage. Thus, short-lived cells can avoid much of the cost of maintaining homeostasis and repairing damage, benefitting from simply being replaced through stem-cell division, with selection removing unfit cells through PCD (Goodell and Rando 2015). This, we suggest, explains the reduced extent of age-related increase in transcriptome variation observed in shorter-lived cells (**Fig. 5B**).

### An evolutionary perspective

How might an individual organism benefit from cells of vastly different life expectancies, like neutrophils, which live for days (McCracken and Allen 2014), versus neurons, which live for decades? There are interesting and potentially important microevolutionary consequences of their differences.

Our results focus primarily on ways that organisms maintain functional long-lived cells, but from an evolutionary perspective, there are ways that short-lived cells might avoid deleterious fitness consequences and ultimately benefit the organism. First, absent major investment in repair and maintenance, we anticipate that short-lived cells age faster than long-lived cells. Despite the potentially rapid accumulation of damage in short-lived cells, the organism can facilitate processes that enhance selective removal of these cells. By selectively removing unfit cells, an organism can thus ensure that this population of rapidly aging cells remains healthy, no matter what the age of the organism. Very long-lived cells, like neurons, have little opportunity for such selective removal and replacement and they instead require investment in intrinsic cellular maintenance. From the organismal perspective, we can think of short cell lifespan as a strategy that minimizes damage within organs while limiting the burden of having to maintain cell function for more than a few days.

Some have further argued that short-lived cells evolve to become very liable to damage and dysfunction, and thus to PCD, ultimately benefiting the organism (Krakauer and Plotkin 2002). This idea mirrors Medawar’s ‘Mutation Accumulation’ theory of aging, where older individuals carry late-acting deleterious alleles (Medawar 1952), albeit not to anyone’s benefit. Following on from Medawar’s model, Williams argued in his ‘Antagonistic Pleiotropy’ theory that genes with beneficial effects early in life would be favored by natural selection, even if they had very deleterious effects at late ages. Any cost of these late-acting deleterious effects is deeply discounted due to the relatively weak force of selection at late age (Williams 1957). Although Williams was thinking about fitness effects of genes on individuals within a population, we can also apply this thinking to the evolution of cells within organisms. If cellular products are beneficial in the short term, but cause problems in the long run, a short-lived cell might be able to explore a greater range of the possible phenotypic space without having to face the long-term costs of those traits. In this sense, short-lived cells might benefit an organism by having fewer antagonistically pleiotropic effects.

### Caveats

There are several limitations to this study worth considering. First, we restricted our analysis to post-mitotic cell types to avoid the confounding effect of mitotic division such as cell self-renewal, differentiation, and senescence, as these cellular processes are associated with stem cells or progenitor cells and might be under different regulatory controls or experience different selection pressures (Brunet et al. 2022). Whether mitotic cells show a similar pattern to the one we observe is an open question. Second, while we did not observe age-related changes in the transcriptomes of short-lived cells when comparing young and old mice, this does not prove that short-lived cells do not experience aging within their short lifetimes. In short-lived cells, the great majority of cells are consistently young, regardless of organismal age (**Fig. 1**). But given that we cannot distinguish young from old cells whose ages differ by just a matter of days, we lack the resolution to observe patterns of transcriptome variation that may vary with cellular age in those cell types. What *is* clear is that when comparing the transcriptome in a population of short-lived cells in young vs. old mice, these mostly young cells show little difference in transcriptome variation across mouse age. Lastly, for simplicity, we assume that the rates of cell turnover do not vary with age, and yet there is evidence that cell turnover can vary with age, and may do so in a cell type-specific manner (reviewed in Tower 2015). We lack information on age-related cellular turnover and so examining its effect on transcriptome variation is not currently possible. However, our results indicate that the effect of cell turnover on the aging transcriptome is clearly predicted by the cellular age distribution.

## Methods

### Mouse transcriptome data

We obtained fluorescence-activated cell sorting-Smart-seq2 profiles of multiple tissues from young (3-month-old) and old (24-month-old) male mice from the *Tabula muris senis* (TMS) database (Zhang et al. 2021). We selected cell types with at least 50 cells in both young and old mice, which gave 52 cell types among 22 organs, resulting in transcriptome profiles from 47,898 total cells. Some cell types are detected in multiple organs (e.g., endothelial cells), and so together the filtered dataset contained 72 combinations of organ:cell type. To account for differences in the number of cells in the two age groups, cell numbers were down-sampled for each organ:cell type separately, so that equal numbers of young and old cells were used. To account for differences in total captured transcript counts across cells, all cells were down-sampled to have an equal number of transcripts in the two age groups for each organ:cell type. After removing genes expressed in less than 10% of cells in each organ:cell type, and cells that express fewer than 100 genes, 35,495 cells remained in our data set.

### Cell type-specific lifespan data

We obtained cell lifespan data, where a cell’s lifespan is defined as the number of days since its last division until its death, from fully differentiated post-mitotic cell types in the available organ:cell type contexts in rodents (Sender and Milo 2021). Twenty one of these organ:cell type combinations corresponded to 39 such combinations in the TMS (**Supplemental Table S1**). All but one organ:cell type have lifespan estimates based on rodents, whereas an estimate for neurons in the nervous system was only available from human studies (Sender and Milo 2021; Spalding et al. 2013, **Supplemental Table S1**). Lifespan estimates in Sender and Milo (2012) involved tracking radiolabeled cells over time, and fitting the data to a model to estimate cellular lifespan. This statistical framework may give lifespan estimates that exceed an animal’s typical longevity (Sender and Milo 2021). Of the 21 organ:cell types in the TMS, 14 have lifespan estimates that were specific to both the cell type and their organ context, such as endothelial cells in brain, heart, or muscle. The remaining 7 cell types lacked context-specific estimates, and so each shares a lifespan estimate across the organs in which they were found. For example, skeletal muscle satellite cells in limbs and in the diaphragm, share the same lifespan as estimated for skeletal muscle satellite cells (Sender and Milo 2021). It is possible that these cells do, in fact, have different lifespans. Our simplifying assumption of common lifespan is a conservative one, as it adds error to our estimates, and so is likely to reduce the magnitude of any real statistical relationship.

### Modeling cell population dynamics

We were able to obtain data on mean cell longevity. However, we recognize that the distribution of cellular ages in an animal of a given age will be a function both of cell turnover rates, and of that individual’s chronological age. Since the maximum cellular age cannot exceed the organism’s age, we used a truncated exponential distribution with an upper bound set by the organismal age. The mean lifespan of a population of cells can be calculated as the first moment of a truncated exponential distribution.

We first transformed cellular lifespan of each cell type in days into a finite survival rate per day (**Supplemental Methods Eq. 1**). We then built a demographic model of cell population dynamics with a two-compartment tissue architecture (**Supplemental Methods, Eqs. 2-4**). Assuming constant cell population size, we derived cell type-specific turnover rate (**Supplemental Methods, Eq. 5**), the cell age distributions (**Supplemental Methods, Eqs. 6-7**), and the analytical solution of the mean cellular age (**Supplemental Methods, Eq. 8**). We plugged in various values of cell type-specific survival rates and the two mouse ages into **Supplemental Methods, Eq. 8** to obtain Δage (**Supplemental Methods, Eq. 9**), or the change in the mean cell age of each cell type within young and old mice shown in **Figure 2**.

### Transcriptome variation analysis

In estimating transcriptome variation, we need to control for the confounding effects of the mean-variance relationship on expression abundance (Eling et al. 2018). To address this issue when evaluating the variation of individual genes in young and old animals, we took a conservative strategy and restricted our analysis to genes whose expression did not change significantly with age, an approach used before in studying the effect of age on immune cells (Martinez-Jimenez et al. 2017). We used a linear model to quantify the age effect on gene expression level using the MAST package in R (Finak et al. 2015). Specifically, we treated all cells of one organ:cell type as samples and performed separate tests for the effect of age on gene expression for each gene.

We removed genes with a P value < 0.1 for a given organ:cell type and then calculated the coefficient of variation (CV) at each age for each of the remaining genes, and tested for effects of age on CV using a linear mixed model with age as a fixed effect and gene as a random effect (*lmer*(*log*(*CV*^2^) ∼ *age* + (1|*gene*)), via the lme4 package in R (Bates et al. 2015). We express the change in transcriptome variability captured by this approach as the coefficient β associated with the effect of age (β_age_).

To analyze the level of transcriptome variability at the cellular level, we estimated the similarity in gene expression profiles among cells using two metrics; Spearman’s rank correlation coefficient (Angelidis et al. 2019) and global coordination level (GCL) (Levy et al. 2020). Spearman’s rank correlation coefficient (ρ) was calculated on the down-sampled expression matrices in all pairwise cell comparisons within each organ:cell type and by age group. We then used 1–ρ as a measure of cell-level transcriptome variation for every cell pair (**Supplemental Fig. S4**). We express the change in transcriptome variability captured by this measure as log2(O/Y), where the median of 1 − ρ in the young sample is Y and that of the old is O.

The second method, GCL, randomly splits genes into two sets and quantifies their mutual dependence via a distance correlation metric between cell samples evaluated on the two gene sets. This metric was shown to capture both linear and non-linear gene correlations and is robust to noise in single-cell data (Vaknin et al. 2021). Highly correlated gene expression patterns among cells give a high GCL, while a low GCL value indicates low levels of similarity in gene expression profiles. We first measured GCL per organ:cell type per age group, which was then transformed into a metric describing the lack of gene coordination, or gene discoordination (1-GCL). Specifically, we randomly split the transcriptome per organ:cell type per age group into halves 100 times and extracted the median value of the resulting 1-GCL values to represent the age-specific discoordination value (**Supplemental Fig. S5**). Similarly, we expressed the change in transcriptome variability captured by these measures as log2(O/Y).

### Cell type-specific gene ontology term expression

We extracted all 12,545 GO terms in the Biological Process (BP) category in mice via the R package org.Mm.eg.db (version 3.14.0) (Marc Carlson 2021). Searching all these GO terms using ‘chaperone’, ‘DNA repair’, and ‘autophagy of mitochondrion’ as the key words, we retained 30 GO terms, 11 of which contained at least seven gene members (**Supplemental Fig. S8**). We calculated a GO term expression for these GO terms for each cell as the proportion of expression abundance among genes in each GO term of the total expression abundance in that cell. Expression levels came from a cell × gene matrix, normalized by the total number of UMIs in each cell and scaled by 10^6^ to counts per million (CPM). In this way, every cell has an expression for each GO term that represents the relative expression of genes in each GO term. We took the mean expressions of cells of a given cell type to represent a cell type-level GO term expression. We calculated the non-parametric correlation between cell lifespan and GO term expression via Kendall’s **τ**.

### Evolutionary constraint analysis

The mode of selection acting on a gene is measured by the nonsynonymous/synonymous substitution rate (*d*_N_/*d*_S_) ratio of mouse-rat orthologous genes. We retrieved *d*_N_/*d*_S_ ratios of mouse-rat orthologous genes from the Ensembl Biomart (v.99) (Yates et al. 2020). Only one-to-one orthologs estimated by Ensembl were included in the study, which gave 14,185 gene pairs. To evaluate *d*_N_/*d*_S_ among cell type-specific genes, we identified cell type-specific genes using the specificity score approach (Sonawane et al. 2017).

For gene *j* in cell type *k*, the specificity score, 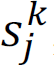, is calculated as:

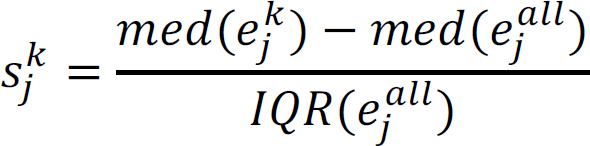

where 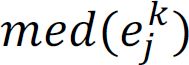 is the median expression level of gene *j* in cell type *k* from a cells × genes matrix, normalized by the total number of UMIs in each cell and scaled by 10^6^ to yield CPM, 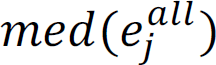 is the median of gene *j* in all cells, and *IQR* is the interquartile range.

To measure *d*_N_/*d*_S_ of the cell type-specific genes in each cell, we took the median *d*_N_/*d*_S_ of the corresponding genes. At each specificity score cutoff, only cells that express at least 100 corresponding cell type-specific genes would be included in the analysis. To represent the evolutionary constraint for each cell type, we took the mean of the *d*_N_/*d*_S_ value of all cells per cell type.

## Data access

This study used previously published datasets (Sender and Milo 2021; Zhang et al. 2021). Cell lifespan data can be accessed from Supplementary Table 5 in Sender and Milo 2021. The TMS data can be accessed from https://figshare.com/articles/dataset/tms_gene_data_rv1/12827615. The code used in this study is available at GitHub (https://github.com/mingwhy/cell.dynamics_aging.asynchrony).

## Acknowledgements

We thank veterinary pathologist Dr. Kim Waggie (University of Washington) for the curation of the alignment of cells in Sender and Milo (2021) (Sender and Milo 2021) with cells in the TMS (Zhang et al. 2021). Cartoons in **Figure 5** were created with BioRender.com. This work was supported by NIH R01 AG063371 (to D. Promislow and S. Pletcher) and Norn Group Impetus Grant (to D. Promislow).

## Author Contributions

M.Y., B.H., and D.P. designed research. M.Y. performed research. M.Y. analyzed data. M.Y., B.H. and D.P. wrote the paper.

