## Supplemental_Material for "Cell age drives asynchronous transcriptome aging"

**This PDF file includes:**

Supplemental Methods

Supplemental Figures S1 to S10

Legend for Supplemental Table S1 and S2

### Supplemental Methods

#### Modeling cell population dynamics

To estimate the age distribution of specific cells, we assume a constant (age-independent) survival rate for a given cell type. We first transformed cellular lifespan of each cell type in days into survival rate per day. Given a finite survival rate per day as $p$, the instantaneous mortality rate $\mu = -ln\ (p)$ (Caswell 1972). Assuming an exponentially distributed survival function, the expected mean lifespan is $T = \frac{-1}{ln\ (p)}$. Inversely, given cellular lifespan, we can calculate finite survival rate as

$$p = exp\left(-\frac{1}{T}\right) \qquad (1)$$

which ranges from 0 to 1.

Using estimates of $p$, we constructed a demographic model describing cell population dynamics, adapted from a standard class-structured life-history model (Caswell 1997), to derive cell type-specific turnover rates, and then working back from cell turnover rates, to determine the cellular age distribution at any organismal age. Considering the simplest case which only contains a progenitor compartment and the downstream fully-differentiated compartment holding cells of different age classes, we aim to derive the distribution of cellular ages in the downstream compartment provided a continuous influx of cells from the progenitor compartment. A life-cycle graph for this model is shown in **Supplemental Figure S1**. For the purposes of this model, we make the following simplifying assumptions. Initially, there are $C$ cells in the cell population at the onset of maturation, and all cellular ages are at age class A=0. At each time unit, three events happen. The progenitor compartment produces $F$ new cells which flow into age class A=0. Existing cells advance from one age class to the next age class with a survival probability of $p$ irrespective of cellular age, or die with a probability of *1-p*. Cell turnover starts at the onset of maturation and is defined as $b = F/C$.

Assuming a discrete time model, the number of cells in age class $i$ at organismal age $t$ is $N(i, t)$, where, $N(i, t)$ can be formulated as

$$N(i,t) = p^i * F, \qquad for\ 0 \le i \le t-1$$

$$N(i,t) = p^i * C,\ for\ i = t \qquad (2)$$

Thus, at organismal age $t$, the total number of cells in a cell population, $S(t)$, is

$$S(t) = \sum_{i=0}^{t} N(i,t) = F + pF + p^2F + \cdots + p^{t-1}F + p^tC$$

$$= \sum_{i=0}^{t-1} p^{t-1} * F + p^t * C$$

$$= \frac{F(1-p^{t-1})}{1-p} + p^t * C \quad (3)$$

We assume that the cell population size $C$ remains constant, and so, dividing both side of equation (3) by $C$, then we have

$$\frac{S(t)}{C} = \frac{F(1-p^{t-1})}{C*(1-p)} + p^t \Leftrightarrow 1 = \frac{F(1-p^{t-1})}{C*(1-p)} + p^t \Leftrightarrow$$

$$1 - p^t = \frac{F(1-p^{t-1})}{C*(1-p)} \Leftrightarrow 1 = \frac{F}{C*(1-p)} \Leftrightarrow$$

$$F = C * (1-p) \quad (4)$$

Thus, under the assumption of constant cell population size, the three parameters must satisfy the above relationship. This makes intuitive sense. In order for the cell population to stay constant, the number of cells lost from the fully differentiated compartment per time unit, $C * (1-p)$, should be equal to the replenishment ($F$) from the progenitor compartment. As a result, the cell turnover rate is

$$b = F/C = (1-p) \quad (5)$$

Next, to get the number of cells in age class *i* at organismal age *t*, we replace $F$ with $C * (1-p)$ in the equation (2) of *N*(*i, t*),

$$N(i,t) = p^i * C * (1-p),\ for\ 0 \le i \le t-1$$

$$N(i,t) = p^i * C,\ for\ i = t \quad (6)$$

The proportion of cells in age class $i$ at organismal age $t$ as $f(k,t)$ is given by dividing equation (6) by $C$ on both sides:

$$f(i,t) = (1-p) * p^i, when\ i = 0, \ldots, t-1$$

$$f(i,t) = p^i, when\ i = t \quad (7)$$

This closed-form analytical expression allows the derivation the mean of the cellular age distribution at organismal age *t*,

$$E(A) = \sum_{i=0}^{t} i * f(i,t) = t * p^t - \frac{t*p^{t+1} - p^{t+1} - t*p^t + p}{p-1} \quad (8)$$

As shown above, E(*A*) is fully determined by cell survival rate, *p*, and the organismal age, *t*. As *p* approaches 1, E(*A*) approaches *t* (i.e., $\lim_{p \to 1} E(A) = t$).

From equation (5), the cell turnover rate is $b = 1 - p = 1 - exp\left(-\frac{1}{T}\right)$. Assuming the onset of maturation in mice at 2 months old (Brust et al. 2015) and using equation (8), we computed the mean cellular age over organismal age for each cell type (**Supplemental Fig. S2**).

At a fixed cell turnover rate $b$, $E(A)$ changes with organismal age $t$. Denoting the mean cell age at 3 months old as E($A$ | $t$ = 3 months) and that at 24 months old as E(A | t = 24 months) , $\Delta_{age}$, or the change in the mean cell age of each cell type within 3-month-old and 24 month-old mice, is calculated as,

$$\Delta_{age} = E(A \mid t = 24 \text{ months}) - E(A \mid t = 3 \text{ months}) \quad (9)$$

### Supplemental Figures

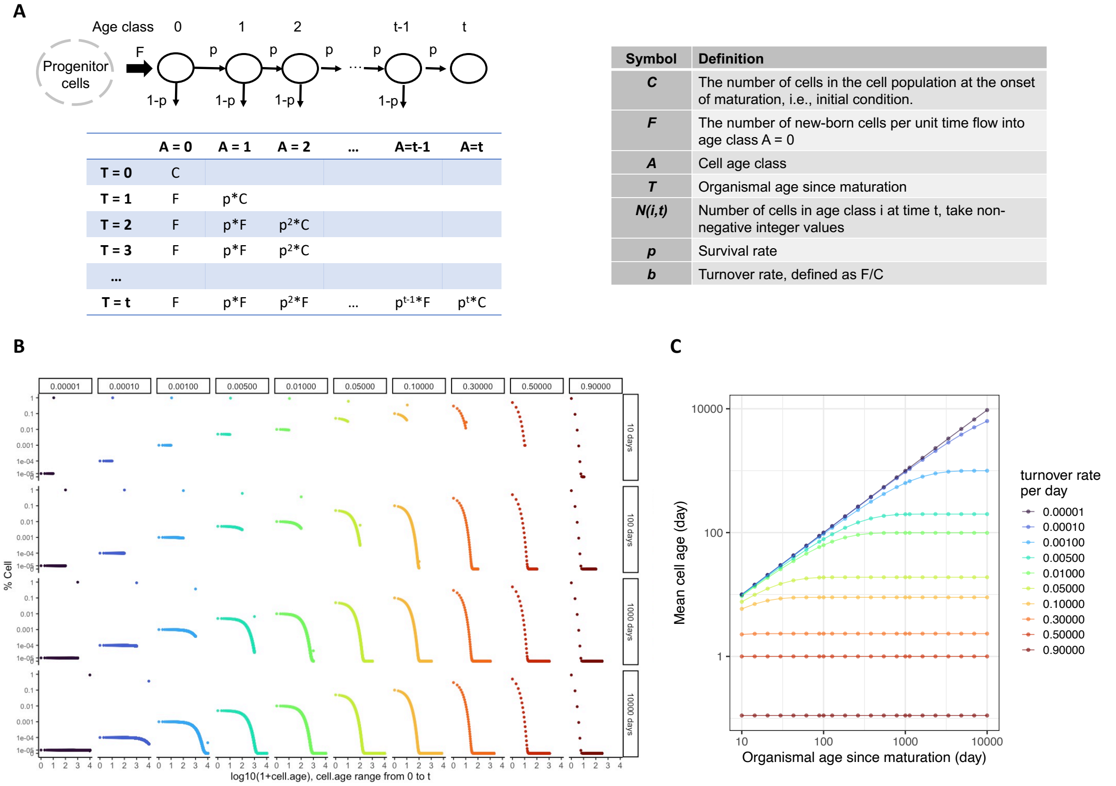


| | A = 0 | A = 1 | A = 2 | ... | A=t-1 | A=t |
|---|---|---|---|---|---|---|
| T = 0 | C | | | | | |
| T = 1 | F | p*C | | | | |
| T = 2 | F | p*F | $p^2$*C | | | |
| T = 3 | F | p*F | $p^2$*C | | | |
| ... | | | | | | |
| T = t | F | p*F | $p^2$*F | ... | $p^{t-1}$*F | $p^t$*C |

| Symbol | Definition |
|---|---|
| *C* | The number of cells in the cell population at the onset of maturation, i.e., initial condition. |
| *F* | The number of new-born cells per unit time flow into age class A = 0 |
| *A* | Cell age class |
| *T* | Organismal age since maturation |
| *N(i,t)* | Number of cells in age class i at time t, take non-negative integer values |
| *p* | Survival rate |
| *b* | Turnover rate, defined as F/C |

**Figure S1. A population dynamic model of cell age distribution.** (**A**) An age-structured life-history model of cell population dynamics. Each black circle represents one age class. Cells in class *i* might die with probability *1-p* or advance into the next age class *i*+1 with probability *p* per time unit. We assume a constant cell population size, with the progenitor compartment producing *F* new cells flowing into age class A = 0 each time unit. The matrix shows the distribution of cells in each age class over organismal ages. (**B**) The cell age distributions at different organismal ages with varying turnover rates, following Equation (7) in the Supplemental Methods. Each column represents one specification of cell turnover rate and each row represents one given organismal age. (**C**) The mean of cellular ages of cell populations at different organismal ages with varying turnover rates. Colors denote different cell turnover rates.

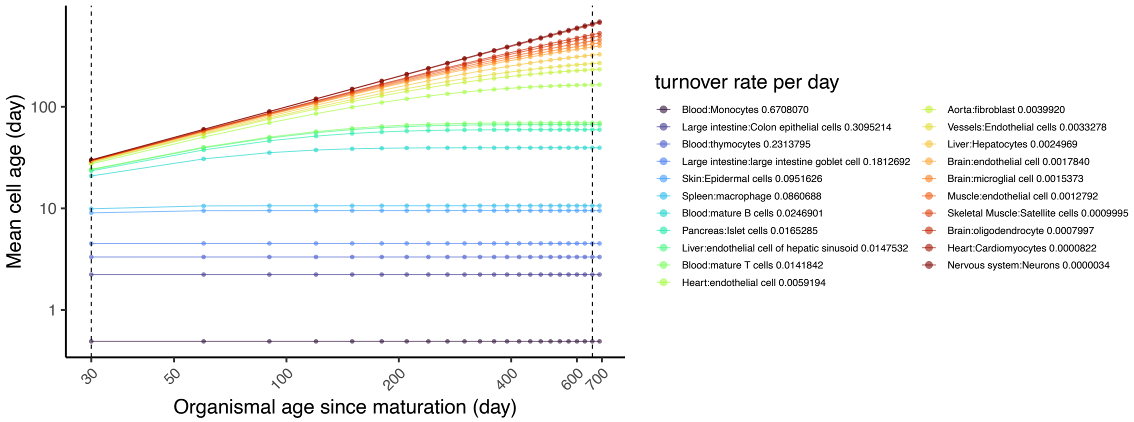


**Figure S2. The effect of organismal age on cell age.** Based on the model presented in the Supplemental Material, this figure shows the relationship between cell type-specific turnover rates and organismal age for specific cell types. Rodent cell-type specific lifespan data are from Sender and Milo (2021). Colors denote different cell types each with its own turnover rate.

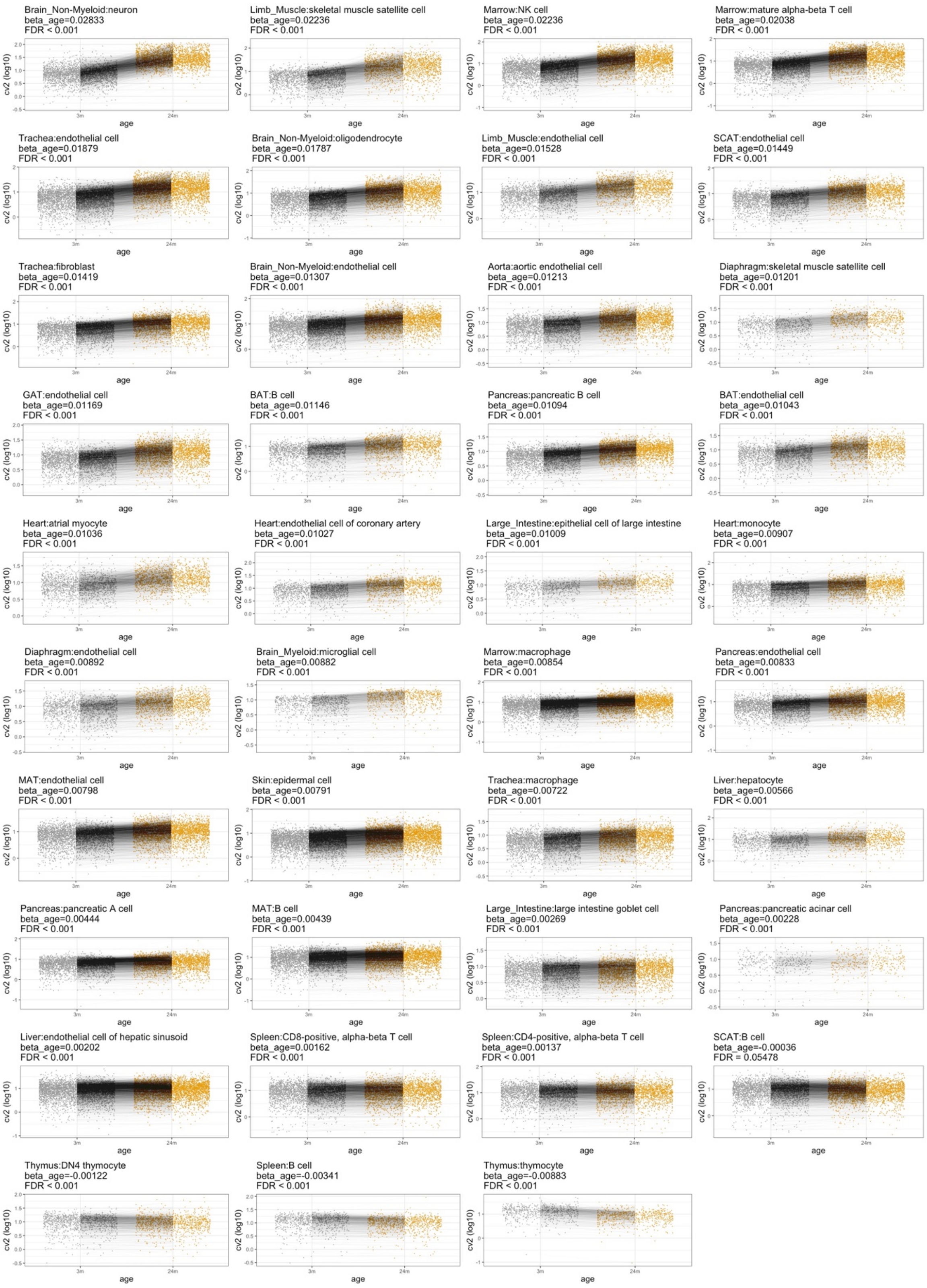


**Figure S3. Effect of age on transcriptome variability**. Each panel represents one organ:cell type combination. In each panel, the squared coefficient of variation ($CV^2$) is plotted for each gene at each age. To avoid an effect driven by a mean-variance correlation, this analysis was limited to genes that did not show age-related changes in mean expression levels (P value > 0.1, Methods in the main text). In each panel, lines connect the same gene measured at the two ages. The number of genes per organ:cell type ranges from 375 to 3962. Age-related changes in gene expression variability are represented by the slope of these lines. For each organ:cell type, the slope ($\beta_{age}$) describing the change in $CV^2$ with age was estimated using a linear mixed model with age as a fixed predictor and gene as a random effect ($\mathrm{lmer}(\log(CV^2)\sim age+(1|gene))$), via the lme4 package in R (Bates et al. 2015).

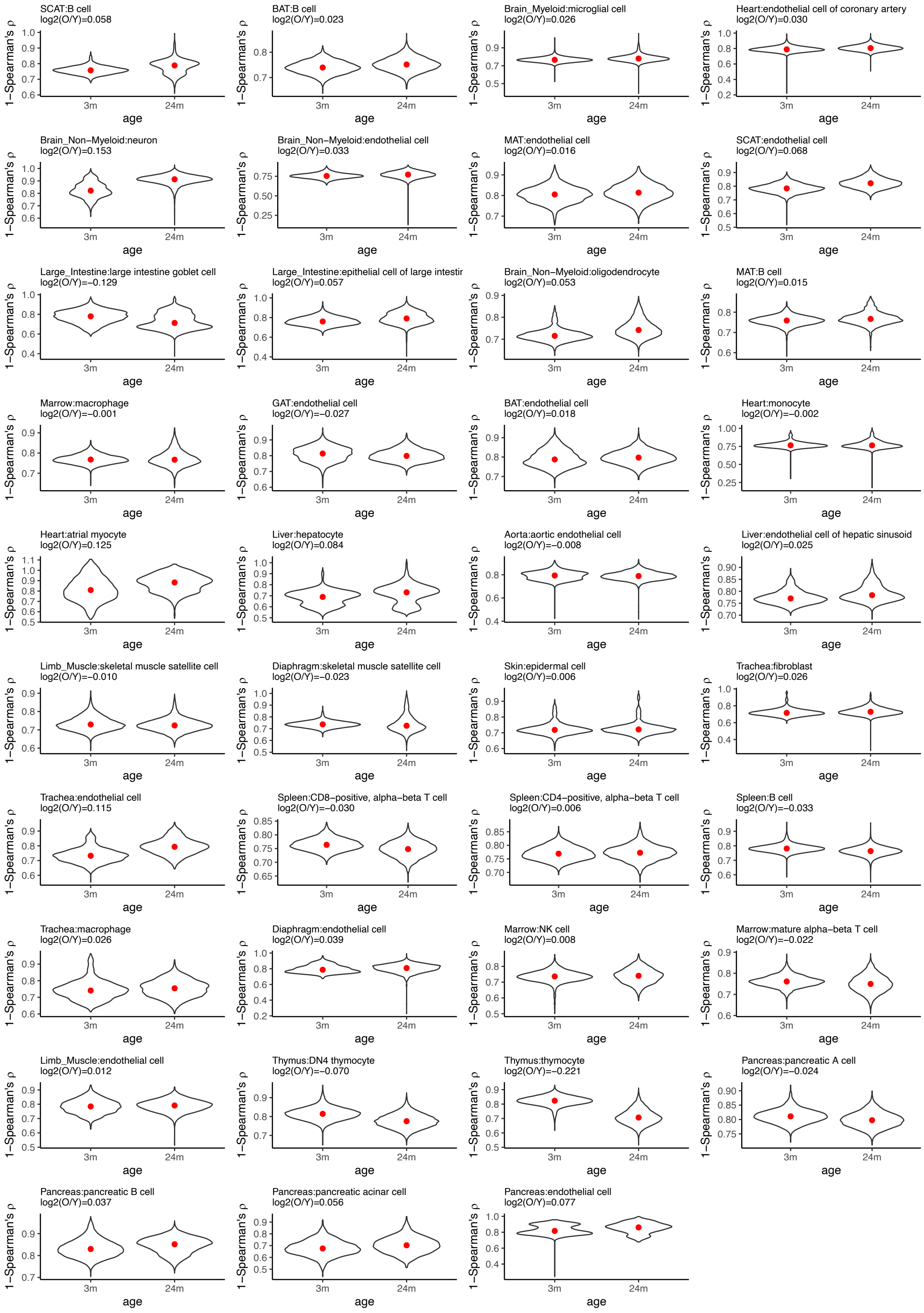
SCAT:B cell
log2(O/Y)=0.058
BAT:B cell
log2(O/Y)=0.023
Brain_Myeloid:microglial cell
log2(O/Y)=0.026
Heart:endothelial cell of coronary artery
log2(O/Y)=0.030
Brain_Non−Myeloid:neuron
log2(O/Y)=0.153
Brain_Non−Myeloid:endothelial cell
log2(O/Y)=0.033
MAT:endothelial cell
log2(O/Y)=0.016
SCAT:endothelial cell
log2(O/Y)=0.068
Large_Intestine:large intestine goblet cell
log2(O/Y)=−0.129
Large_Intestine:epithelial cell of large intestir
log2(O/Y)=0.057
Brain_Non−Myeloid:oligodendrocyte
log2(O/Y)=0.053
MAT:B cell
log2(O/Y)=0.015
Marrow:macrophage
log2(O/Y)=−0.001
GAT:endothelial cell
log2(O/Y)=−0.027
BAT:endothelial cell
log2(O/Y)=0.018
Heart:monocyte
log2(O/Y)=−0.002
Heart:atrial myocyte
log2(O/Y)=0.125
Liver:hepatocyte
log2(O/Y)=0.084
Aorta:aortic endothelial cell
log2(O/Y)=−0.008
Liver:endothelial cell of hepatic sinusoid
log2(O/Y)=0.025
Limb_Muscle:skeletal muscle satellite cell
log2(O/Y)=−0.010
Diaphragm:skeletal muscle satellite cell
log2(O/Y)=−0.023
Skin:epidermal cell
log2(O/Y)=0.006
Trachea:fibroblast
log2(O/Y)=0.026
Trachea:endothelial cell
log2(O/Y)=0.115
Spleen:CD8−positive, alpha−beta T cell
log2(O/Y)=−0.030
Spleen:CD4−positive, alpha−beta T cell
log2(O/Y)=0.006
Spleen:B cell
log2(O/Y)=−0.033
Trachea:macrophage
log2(O/Y)=0.026
Diaphragm:endothelial cell
log2(O/Y)=0.039
Marrow:NK cell
log2(O/Y)=0.008
Marrow:mature alpha−beta T cell
log2(O/Y)=−0.022
Limb_Muscle:endothelial cell
log2(O/Y)=0.012
Thymus:DN4 thymocyte
log2(O/Y)=−0.070
Thymus:thymocyte
log2(O/Y)=−0.221
Pancreas:pancreatic A cell
log2(O/Y)=−0.024
Pancreas:pancreatic B cell
log2(O/Y)=0.037
Pancreas:pancreatic acinar cell
log2(O/Y)=0.056
Pancreas:endothelial cell
log2(O/Y)=0.077
1−Spearman's ρ
age
3m
24m

**Figure S4. Effect of age transcriptome variability, as measured by cell-cell correlations.** Cell-to-cell correlation analyses were performed among cells of the same organ:cell type at the same age group, using Spearman's rank correlation coefficients ($\rho$) as the measure of similarity between cells. Each $\rho$ value corresponds to a pair of cells. The number of cells per organ:cell type per age group ranges from 55 to 2253. The y-axis shows 1- $\rho$ and red dots denote the median value. The age effect is measured as log2(O/Y), where the median of $1 - \rho$ in the young sample is Y and that of the old is O. Larger values of O compared with Y indicate larger cell-to-cell distances.

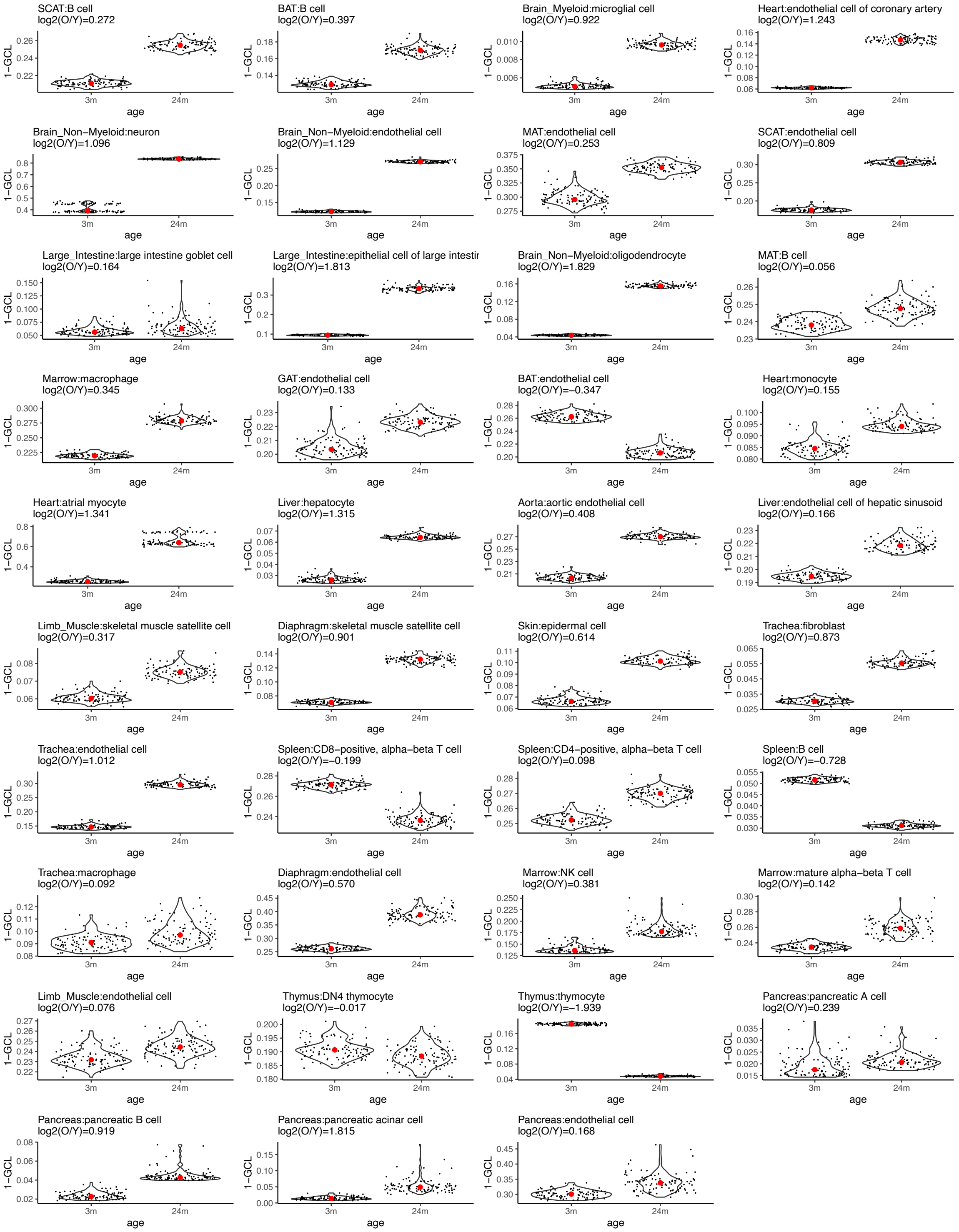


**Figure S5. Effect of age on transcriptome variability as measured by GCL.** The Global Coordination Level (GCL) metric, in which the similarity among cells is assessed, based on their distance in gene expression space within subsets (samples) of genes, has been used to measure similarity among cells (Levy et al. 2020). The y-axis shows the 1-GCL value, which increases with an increase in gene expression divergence. Each dot represents the distance between cells, measured by the similarity of expression matrices, made from the same organ:cell type, when divided in half using random halves of genes expressed in each organ:cell type (Levy et al. 2020). There are 100 such random samples for each organ:cell type at each age group. The red dot inside each violin plot denotes the median value, and the age effect is measured as log2(O/Y), where the median of 1-GCL in the young sample is Y and that of the old is O.

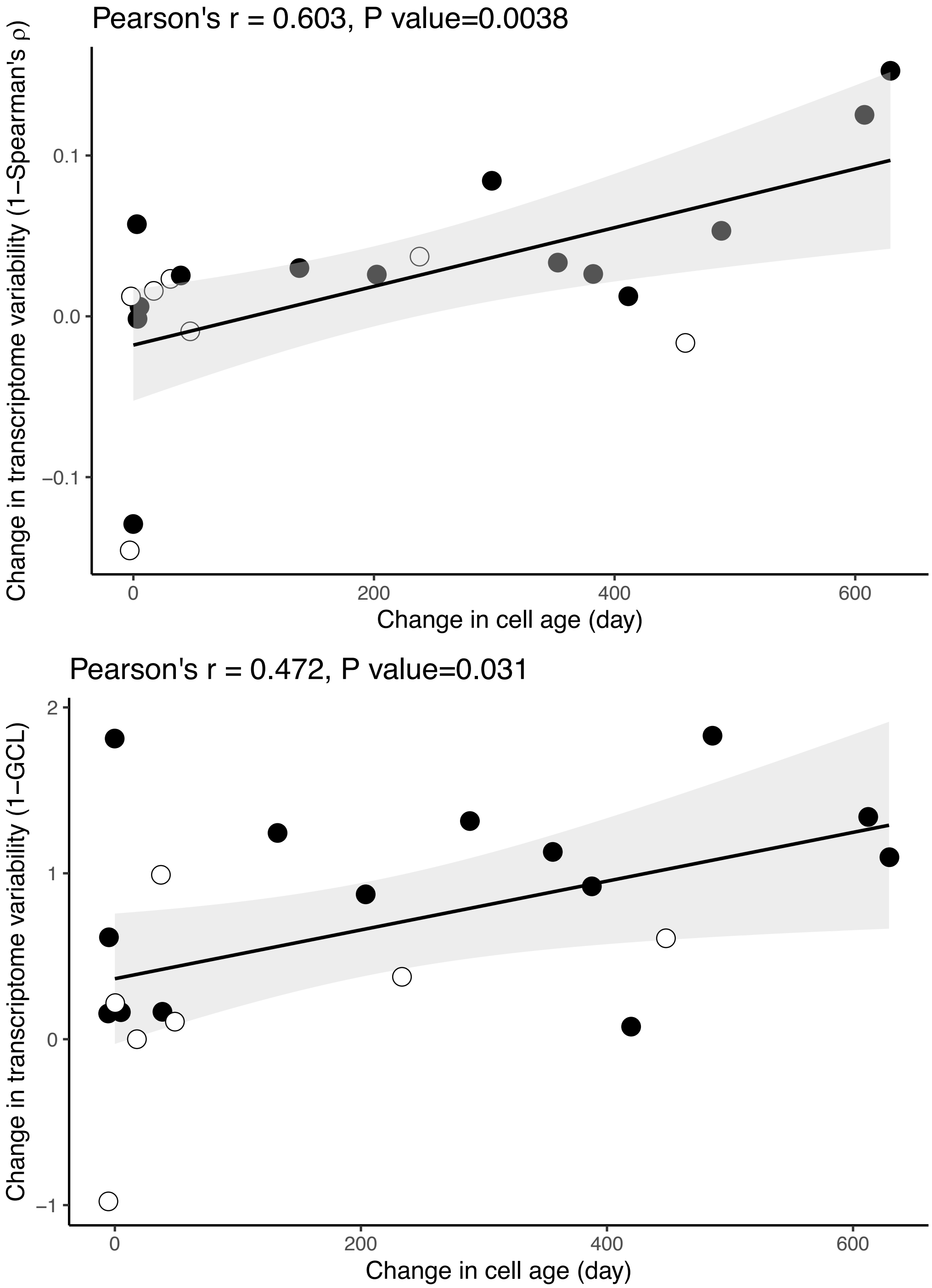


**Figure S6. Alternative metrics for the effect of cell age on transcriptome variability.** The two figures above illustrate the relationship between age-related increases in transcriptome variability and cell age, using two approaches, Spearman rank correlations, and GCL (see Methods). The black line shows an ordinary least squares regression, with shading indicating 95% confidence intervals. Filled circles indicate the 14 cell types with cell- and organ-specific lifespan data, and open circles the 7 cell types without organ-specific lifespan estimates, and so their $\beta_{\text{age}}$ is the mean from all organs in which they are detected (see **Methods**).

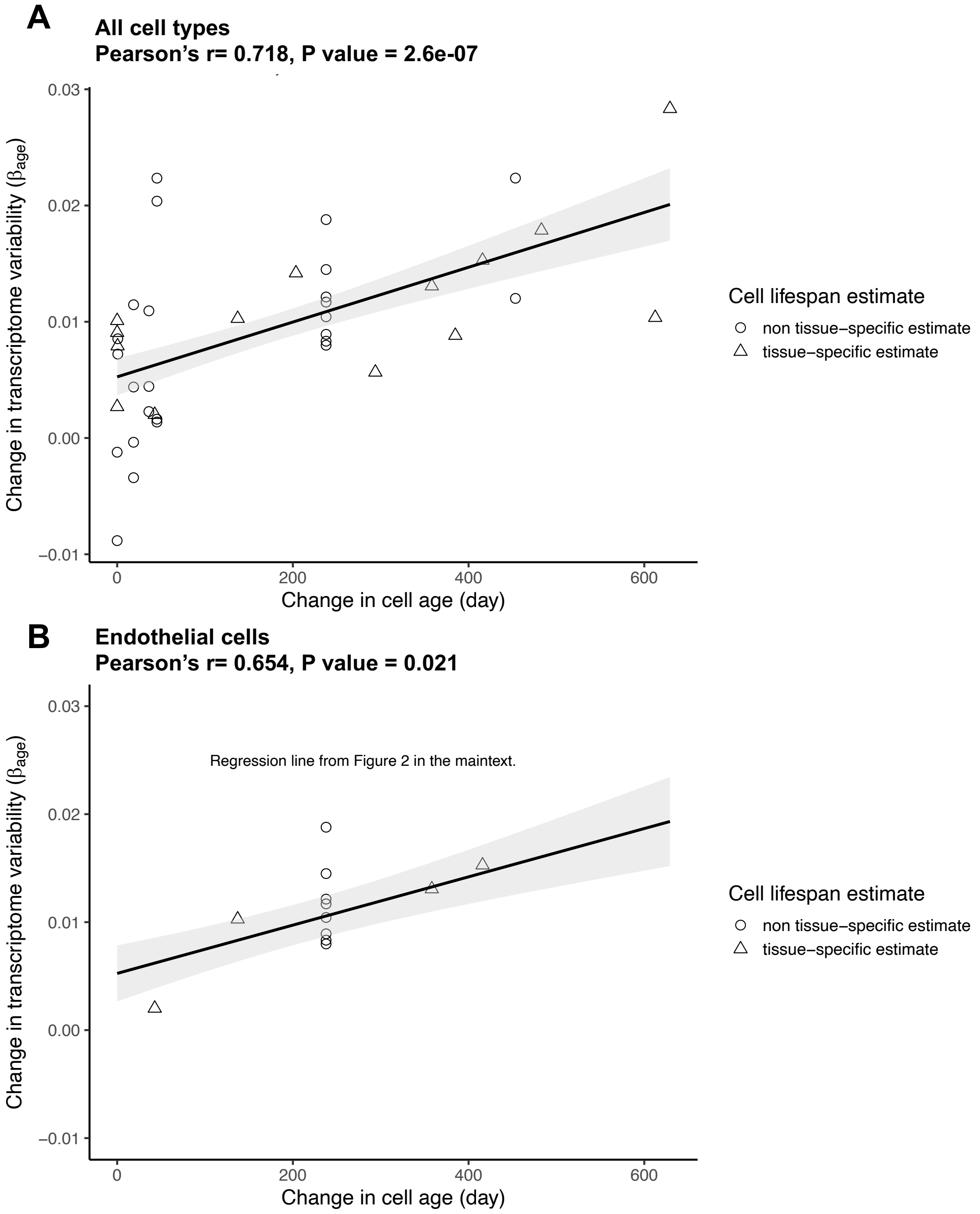


**Figure S7. Transcriptome variability by tissue may be explained by tissue-specific effects on cell turnover. (A)** The change in transcriptome variability with age ($\beta_{age}$) across organ:cell types, where point shapes denote cell types with (triangles) or without (circles) tissue-specific lifespan estimates. **(B)** Among the 12 tissues that had transcriptome data for endothelial cells, there are four with tissue-specific estimates of cell lifespan (triangles), from which we derived four separate cellular age distributions to get the change in cellular age ($\Delta_{age}$). These tissue-specific $\beta_{age}$ and $\Delta_{age}$ values are shown in comparison to the regression of all data (black line taken from **Fig. 2** in the main text).

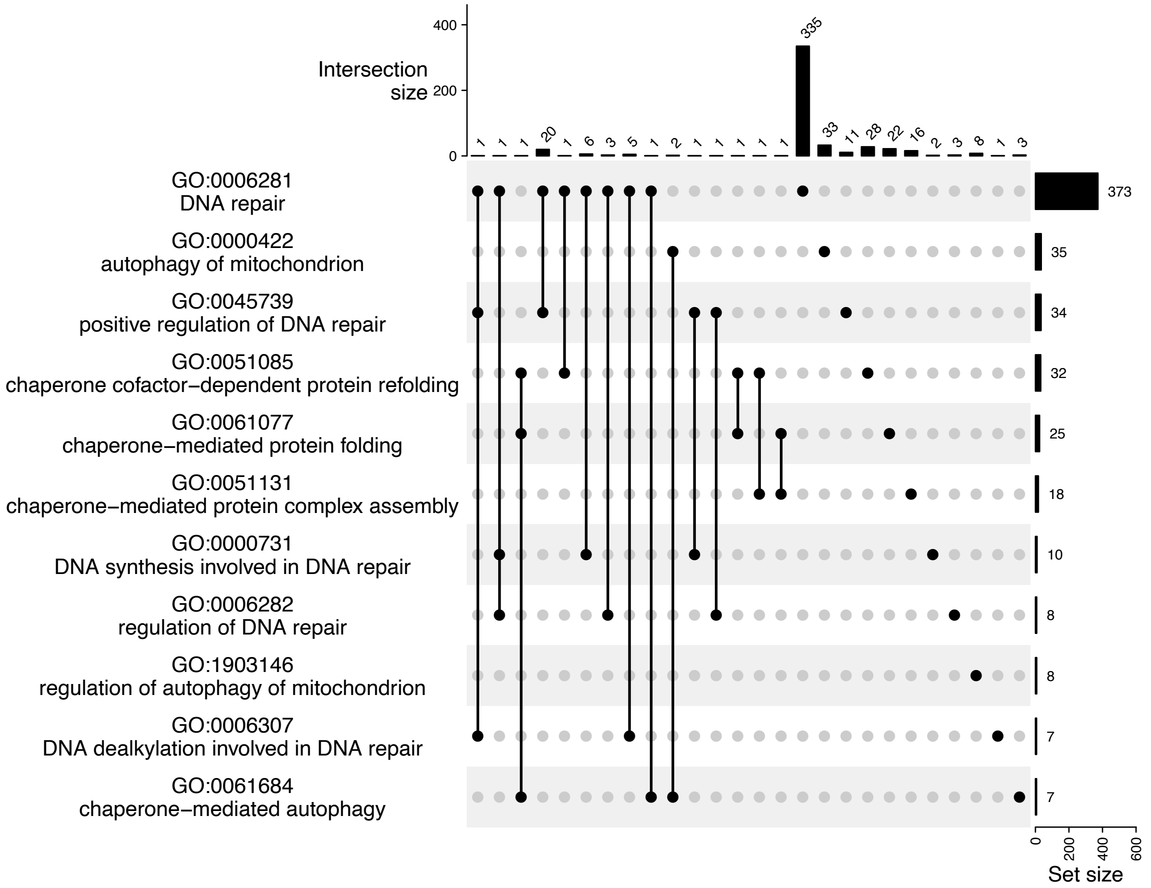


**Figure S8. Intersection between the Gene Ontology (GO) biological process terms related to 'chaperone', 'DNA repair', and 'mitophagy'.** Each GO term contains at least seven gene members shown as the horizontal bars (Set size). The top margin shows the number of genes that intersect between the GO terms. We used gene member expression abundance to calculate GO term expression and limited this analysis to non-redundant terms; for the 11 GO terms, there were at most 20 intersecting genes shared between terms, and nine of the terms shared fewer than six genes.

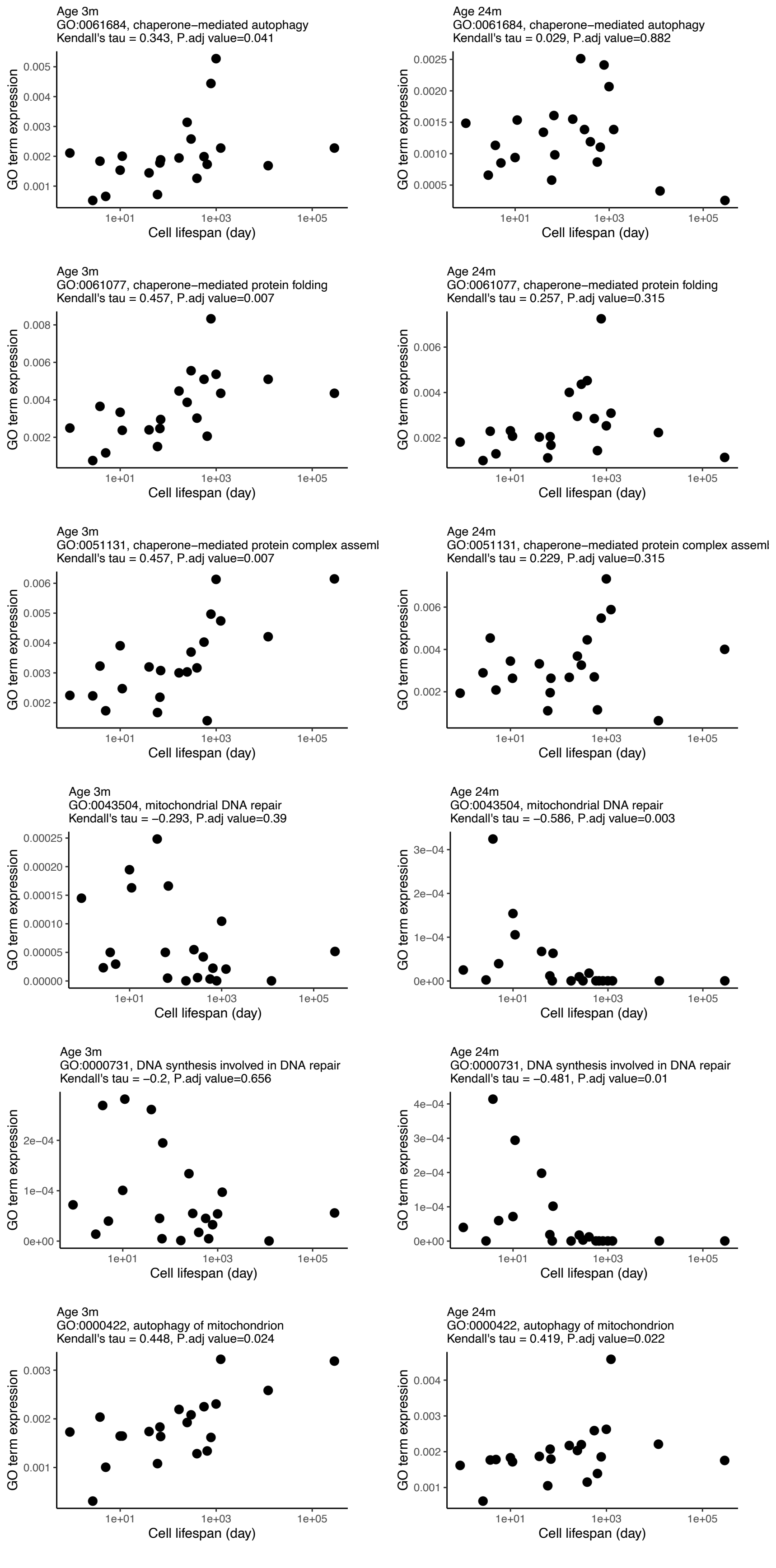
Age 3m
GO:0061684, chaperone−mediated autophagy
Kendall's tau = 0.343, P.adj value=0.041
Age 24m
GO:0061684, chaperone−mediated autophagy
Kendall's tau = 0.029, P.adj value=0.882
Age 3m
GO:0061077, chaperone−mediated protein folding
Kendall's tau = 0.457, P.adj value=0.007
Age 24m
GO:0061077, chaperone−mediated protein folding
Kendall's tau = 0.257, P.adj value=0.315
Age 3m
GO:0051131, chaperone−mediated protein complex asseml
Kendall's tau = 0.457, P.adj value=0.007
Age 24m
GO:0051131, chaperone−mediated protein complex asseml
Kendall's tau = 0.229, P.adj value=0.315
Age 3m
GO:0043504, mitochondrial DNA repair
Kendall's tau = −0.293, P.adj value=0.39
Age 24m
GO:0043504, mitochondrial DNA repair
Kendall's tau = −0.586, P.adj value=0.003
Age 3m
GO:0000731, DNA synthesis involved in DNA repair
Kendall's tau = −0.2, P.adj value=0.656
Age 24m
GO:0000731, DNA synthesis involved in DNA repair
Kendall's tau = −0.481, P.adj value=0.01
Age 3m
GO:0000422, autophagy of mitochondrion
Kendall's tau = 0.448, P.adj value=0.024
Age 24m
GO:0000422, autophagy of mitochondrion
Kendall's tau = 0.419, P.adj value=0.022
GO term expression
Cell lifespan (day)
1e+01
1e+03
1e+05

**Figure S9. The relationship between GO term expression and cell lifespan in mice**. Each panel shows one GO term. The left columns denote results from 3-month old mice and the right from 24-month old mice. Each dot shows one cell type, and correlation was tested using non-parametric Kendall's tau. P values were corrected for multiple comparisons using the FDR method.

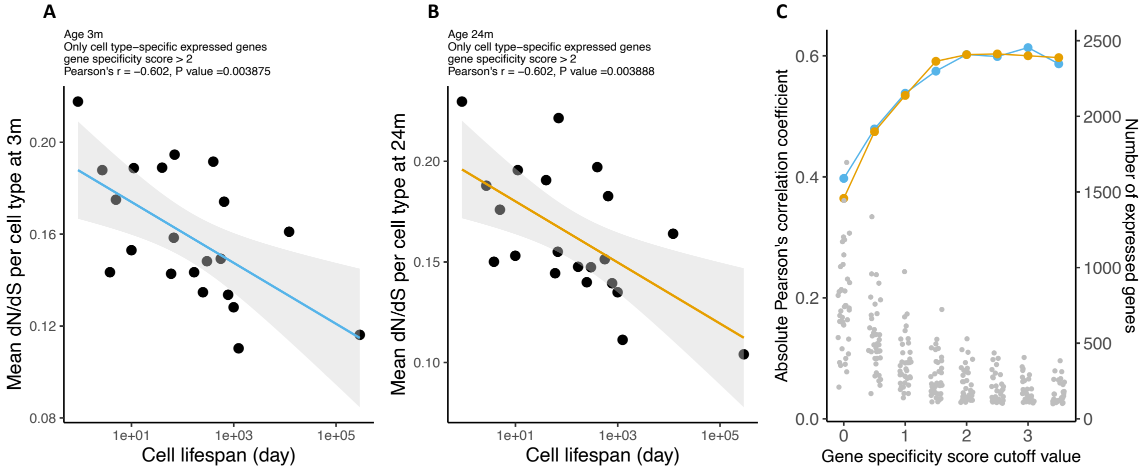


**Figure S10. After removing genes under positive selection, short-lived cells still show relaxed evolutionary constraint.** The relaxation of evolutionary constraint ($d_N/d_S$) of genes expressed in 21 cell types in young (panel **A**) and old (panel **B**) mice is negatively correlated with cell lifespan. Smaller $d_N/d_S$ values connote greater evolutionary constraint. The black line shows the ordinary least squares regression with shading of 95% confidence intervals. The analysis was performed with 14,109 genes after excluding 76 genes whose $d_N/d_S > 1$. **(C)** Correlation between $d_N/d_S$ and cell lifespan for transcriptomes of increasing cell-type specificity. The blue (age 3m) and yellow (age 24 m) dots correspond to the left-side axis, and show the correlation strength. The grey dots correspond to the right-side axis and show the number of expressed genes per cell type at each level of specificity.

**Supplemental Table S1**. Cell types in the *Tabula Muris Senis* **(**TMS) dataset (Zhang et al. 2021) and their respective cellular lifespan estimations in rodents from the public database compiled from the Sender and Milo (2021) study (Sender and Milo 2021).

**Supplemental Table S2**. Change in cellular age in young and old mice and change in transcriptome variability measured by different metrics.
